# NK cells integrate signals over large areas when building immune synapses but require local stimuli for degranulation

**DOI:** 10.1101/2020.07.29.224618

**Authors:** Quentin Verron, Elin Forslund, Ludwig Brandt, Mattias Leino, Thomas W. Frisk, Per E. Olofsson, Björn Önfelt

**Author notes:** **Summary** Using micro-arrays of artificial immune synapses, we demonstrate that the spatial distribution of activating ligands at the NK cell immune synapse can influence the stability and cytotoxic outcome of the synapse, suggesting a new potential escape mechanism for target cells.

## Abstract

Immune synapses are large-scale, transient molecular assemblies that serve as platforms for antigen presentation to B and T cells, and target recognition by cytotoxic T cells and natural killer (NK) cells. The formation of an immune synapse is a tightly regulated, stepwise process where the cytoskeleton, cell-surface receptors and signaling proteins rearrange into supramolecular activation clusters (SMACs). Here we use a reductionist system of microcontact-printed artificial immune synapses (AIS) shaped as hallmark SMAC structures to show that the spatial distribution of activating ligands influences the formation, stability and outcome of NK cell synapses. Organizing ligands into donut-shaped AIS resulted in fewer long-lasting, symmetrical synapses compared to dot-shaped AIS. NK cells spreading evenly over either AIS exhibited similar arrangement of the lytic machinery, however degranulation was only possible in regions allowing local signaling. Our results demonstrate that the macroscopic organization of ligands in the synapse can affect its outcome, which could be exploited by target cells as an escape mechanism.

## Introduction

Natural killer (NK) cells are innate cytolytic immune cells that are crucial for host defense against viral pathogens and cancer. NK cell surveillance relies on recognition of potential target cells, mediated by ligation of activating and inhibitory receptors expressed at the NK cell surface. This process is facilitated by the formation of a tight intercellular contact between the NK cell and its target, *i.e.* an immune synapse. At this interface, NK cell receptors and target cell ligands have been shown to segregate into separate spatial domains called supramolecular activation clusters (SMACs) (Davis et al., 1999). Initial studies of immune synapse formation in T cells and NK cells described a central (c-) SMAC concentrating activating signals, encircled by a peripheral (p-) SMAC promoting adhesion, which in turn is surrounded by a distal (d-) SMAC containing proteins excluded from the immune synapse (Davis, 2002; Grakoui et al., 1999; Monks et al., 1998; Vyas et al., 2001). Although subsequent work has shown that the picture is complex (Almeida & Davis, 2006; Orange et al., 2003; Vyas et al., 2002), NK cell receptors involved in the immune synapse are commonly observed to distribute across the central part of the contact area (corresponding to the cSMAC) or in a peripheral ring structure (corresponding to the pSMAC) (Davis et al., 1999; Orange et al., 2003). Thus, these two distribution patterns are hallmarks for the NK cell immune synapse.

Once the NK cell has engaged in an immune synapse with its potential target, the balance between activating and inhibitory signals determines the outcome of the interaction (reviewed in Sivori et al., 2019). If activating signals dominate, an enclosed cleft is formed in the central parts of the synapse where cytolytic molecules including perforin and granzymes can be released from specialized intracellular vesicles, causing target cell death (reviewed in Orange, 2008). The stepwise formation of this cytotoxic immune synapse involves recognition and adhesion to the target cell, minus-ended movement of lytic granules along microtubules until convergence at the microtubule-organizing center (MTOC), reorganization of the actin cytoskeleton at the synapse, receptor- and ion channel-mediated signaling, polarization of the MTOC and lytic granules towards the intercellular contact, followed by fusion of the lytic granules with the NK cell membrane resulting in the directed secretion of their cytotoxic content into the synaptic cleft (Mace et al., 2014; Mentlik et al., 2009; Orange, 2008). As for cytotoxic T cells, calcium signaling, via entry of extracellular calcium, is crucial for NK cell degranulation (Leibson et al., 1990; Maul-Pavicic et al., 2011; Takayama & Sitkovsky, 1987).

These multiple steps towards cytotoxicity have been shown to be differently driven by individual receptor engagement (Brown et al., 2012; Bryceson et al., 2005). Ligation of the integrin LFA-1 (CD11a/CD18) on the NK cell facilitates adhesion to the target cell and initiates the first steps of immune synapse formation including F-actin polymerization (Barber et al., 2004; Mace et al., 2014; Mentlik et al., 2009). Engagement of LFA-1 alone or in combination with inhibitory receptors results in asymmetric spreading of the NK cell and resumption of migration, whereas its association with an activating signal leads to the formation of a stable, symmetrical synapse (Culley et al., 2009). Furthermore, ligation of LFA-1 has been shown to stimulate polarization but not degranulation of lytic granules, while engagement of the NK cell activating Fc-receptor CD16 induces degranulation but not polarization of lytic granules (Bryceson et al., 2005). Ligation of both LFA-1 and CD16 leads to polarization of the lytic machinery and degranulation in the direction of the target cell (Bryceson et al., 2005).

Recent studies have demonstrated that signaling at the immune synapse is regulated not only by the nature of receptors involved but also by their spatial distribution. As described for receptors engaged in the immune synapses formed by T cells (Bunnell et al., 2002; Kaizuka et al., 2007; Seminario & Bunnell, 2008; Varma et al., 2006; Yokosuka et al., 2005), also NK cell receptors have been shown to form microclusters at the synapse. These include both inhibitory and activating receptors, e.g. NKG2A and NKG2D (Culley et al., 2009), the natural cytotoxicity receptor NKp46 (Hadad et al., 2015) and the Fc-receptor CD16 (Keydar et al., 2018). Signaling from these microclusters can be processed locally or in a cumulative fashion where the NK cell is able to integrate signals from spatially separated ligands (Culley et al., 2009). Modifying the local distribution of NK cell ligands by altering their segregation in separate domains (Martinez et al., 2011) or forcing T cell receptor arrangement into defined static structures (Doh & Irvine, 2006) modulates lymphocyte activation and reorganization of molecular assemblies in the cell.

Here we used microcontact printing to create arrays of artificial immune synapses (AIS) in order to investigate the consequence of ligand distribution on NK cell function. Combining NK cell adhesion via LFA-1 and activation via CD16, receptors were stimulated with typical immune synapse structures in the shape of even disks (“dots”) or rings (“donuts”). Time-lapse imaging of resting NK cells interacting with AIS revealed that the spatial distribution of ligands influenced the formation of the synapse as shown by more frequent transient, partial contacts on donut-shaped AIS. These results show that the NK cell response is regulated not only by the type and abundance of stimuli but also by their spatial distribution. NK cells that showed a strong calcium response often spread symmetrically to cover the whole AIS, regardless of the shape of the AIS. These complete contacts also exhibited similar organization of plasma membrane at the AIS-interface and axial and lateral positioning of the MTOC, indicating that NK cells integrate signaling from spatially separated stimuli when building the IS. However, degranulation was only observed in regions where there were local stimuli from ligands. This suggests that NK cell cytotoxicity is regulated by signal integration over a large area for establishment of an immune synapse and by local stimuli for execution of degranulation.

## Results

### Ligation of LFA-1 alone promotes adhesion and motility while combined engagement with CD16 gives a stop signal that leads to symmetric spreading

To mimic human NK cell immune synapses, microcontact printing (Kumar & Whitesides, 1993) was used to pattern antibodies stimulating the NK cell receptors LFA-1 and CD16 in arrays of dot- or donut-shaped, cell-sized artificial immune synapses (AIS) onto glass substrates suitable for high-resolution, live-cell fluorescence microscopy (Figure 1).

**Figure 1.**
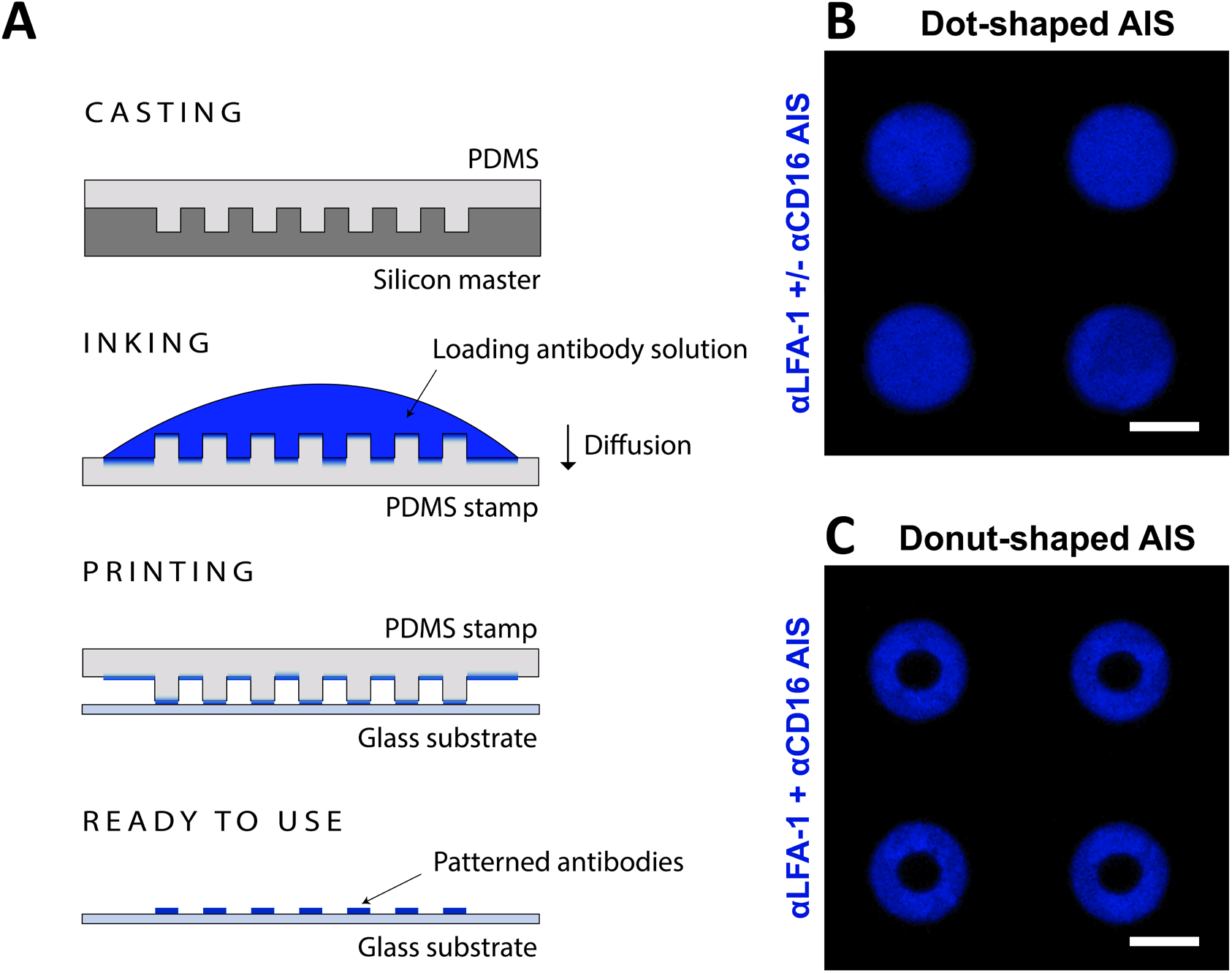
Antibodies patterned into AIS using microcontact printing. **(A)** Schematic representation of the microcontact printing process. Poly(dimethylsiloxane) (PDMS) was molded on a microstructured silicon master (casting). The PDMS was peeled off from the master, loaded with an antibody solution (inking) and briefly dried. The stamp was applied to a poly-L-lysine-coated glass surface, leaving an array of AIS that was used as substrate for imaging live or fixed NK cells. **(B-C)** Example fluorescence images of AIS in dot and donut shapes (C). The loading antibody solution was supplemented with 10 μg/mL BSA labelled with Alexa Fluor 555 (here shown in blue) for visualization. Scale bars: 10 μm.

The dynamic responses of NK cells interacting with dot-shaped AIS stimulating LFA-1 and CD16 were studied by live cell imaging. Consistent with previous observations (Culley et al., 2009; Smith et al., 2003), we found that ligation of LFA-1 alone triggered the initiation of an immune synapse but in the absence of further activating signals the cells remained elongated and asymmetrical (Figure 2A). Combining engagement of αLFA-1 with αCD16 resulted in NK cells stopping and spreading out symmetrically across the AIS to assume morphologies resembling mature activating immune synapses (Figure 2B). This behavior manifested in larger spreading area (Figure 2C) and increased roundness (Figure 2D) on AIS with both αLFA-1 and αCD16. Also, there were marked differences in migration behavior. NK cells migrated faster on arrays of AIS with only αLFA-1 compared to AIS with αLFA-1 and αCD16 (Figure 2E). Using a previously developed method based on detecting transient changes in migration behavior for individual cells, we divided NK cell migration tracks on AIS into periods of arrest, random or directed movement (Huet et al., 2006; Khorshidi et al., 2011). NK cells on arrays of αLFA-1 + αCD16 AIS spent significantly more time in transient migration arrest periods (TMAPs, Figure 2F-H). Directed movement was only rarely observed on AIS of either composition. These differences in cell shape, spreading area and migration behavior were not unique to arrays of AIS as similar differences were observed also on surfaces evenly coated with αLFA-1 alone or a mix of αLFA-1 and αCD16 (Supporting Figure S1). Slightly higher migration speeds, associated with longer periods of directed migration, were measured on evenly coated surfaces, which could be due to the cells having access to a continuous layer of protein that is not present on arrays of AIS.

**Figure 2.**
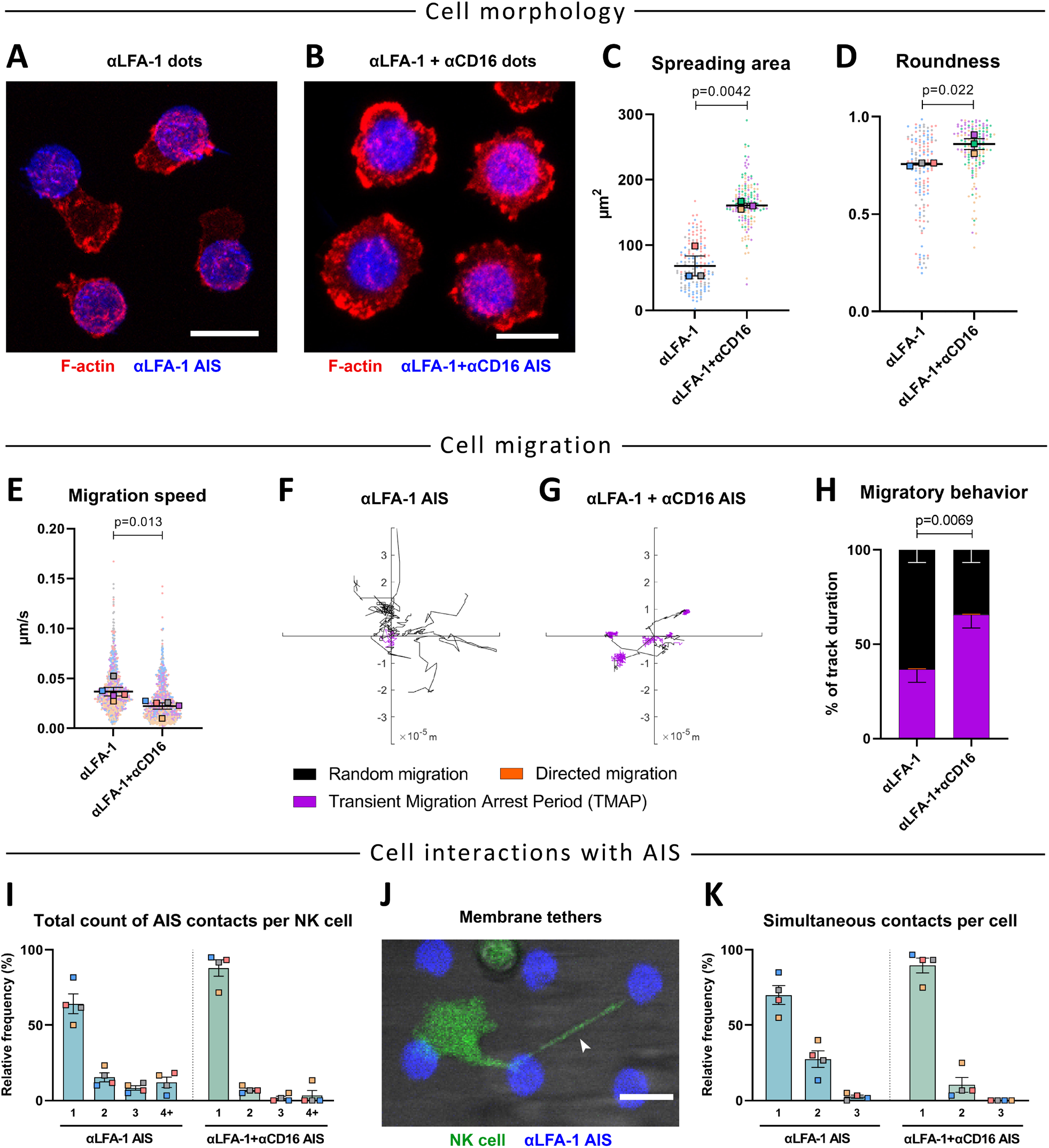
Interaction of NK cells with αLFA-1 AIS leads to a migratory, exploratory phenotype, while combined ligation of LFA-1 and CD16 induces cell arrest and spreading. (A-B) Representative fluorescence images of NK cells interacting with AIS of αLFA-1 (A) or αLFA-1 + αCD16 (B). NK cells were stained for f-actin (red) and maximum intensity projection images were generated from confocal z-stacks. Spreading area of NK cells on either AIS, measured as the footprint of the NK cell f-actin network in the imaging plane of the AIS. **(D)** Roundness of NK cells interacting with either AIS. Data in (C, D) are from 3 independent experiments, N=50 per condition and experiment. (E-H) NK cells were stained with Calcein Green and followed by time-lapse microscopy during their interaction with AIS. **(E)** NK cell migration speed on either AIS. **(F-G)** Example NK migration tracks that have been divided into three different modes of migration; arrest (TMAP, magenta), random (black) and directed migration (orange). **(H)** Average percent of time spent in the different modes of migration on either AIS. The P-value was calculated by comparing the TMAP fraction between groups. Data in (E, H) are from 5 independent experiments, N=150 per condition and experiment. **(I)** Total count of AIS encountered by individual NK cells over the course of the assay (240 min). **(J)** Example fluorescence image of a Calcein Green-stained NK cell (green) in contact with two AIS simultaneously using membrane tethers (white arrow). **(K)** Maximum number of AIS contacted simultaneously by individual NK cells. Data in (I, K) are from 4 independent experiments, N=60 per condition and experiment. Scale bars: 10 μm.

The increased migration on AIS composed of αLFA-1 alone resulted in individual NK cells contacting several AIS on these arrays. On αLFA-1 AIS arrays, about 35% of the NK cells contacted multiple AIS during the 240 min assay while the corresponding number for αLFA-1 + αCD16 AIS was only 12% (Figure 2I). Interestingly, on αLFA-1 AIS NK cells often appeared anchored to a print at one end of the cell while scanning the surroundings with the other end, giving them an elongated shape (as measured in Figure 2D). During this process membrane nanotubes, resembling those that have been observed between NK and target cells (Chauveau et al., 2010; Önfelt et al., 2004) were often formed, facilitating adhesion to one AIS while migrating and binding to additional AIS (Figure 2J). These membrane nanotubes could connect one NK cell with up to three different AIS simultaneously. Similar behavior was not observed on αLFA-1 + αCD16 AIS where the maximum number of simultaneous contacts was 2 (Figure 2K). When AIS were printed with a short center-to-center distance (≤ 30 μm), we observed that some cells assumed an elongated shape, forming stable contacts with two prints at the same time, with the main part of the cell body oscillating between the prints (Supporting Movie 1). These results show that signaling through LFA-1 triggers synapse initiation but allows for continued NK cell motility, that can be balanced by spreading and stopping induced by CD16.

### The spatial distribution of αLFA-1 and αCD16 influences the stability of cell contact by regulating the dynamics of NK cell spreading across the AIS

To investigate if NK cell responses were affected by the spatial distribution of ligands, NK cells were imaged on either dot- or donut-shaped AIS containing equal parts of αLFA-1 and αCD16. NK cells formed significantly longer contacts on dot-shaped AIS with 90% of contacts lasting two hours or longer, while the corresponding fraction was 57% for NK cells adhering to donut-shaped AIS. The mean contact times measured were 199 min for dot-shaped AIS and 129 min for donut-shaped AIS (Figure 3A). These results could be explained by differences in how the NK cell interacted with the AIS. On dots, NK cells often formed contact with the print from one side or directly at the center and spread out symmetrically over the printed area to eventually cover the entire AIS (Figure 3B). On donut AIS contacts were initiated on the antibody ring but due to the void of activating ligands in the central region, gradual symmetrical spreading across the AIS was hindered. Instead NK cells often moved along one side of the print (Supporting Movie 2) or even both sides without spreading across the center (Figure 3C). NK interactions with AIS were thus classified as either complete contacts, if the NK cell spread out to eventually cover the entire print (Supporting Movie 3), or as partial contacts if not. Complete contacts were significantly more frequent on dot-shaped AIS (75%) compared to donut-shaped AIS (46%) (Figure 3D). NK cells forming complete contacts on either dot- or donut-shaped AIS showed a similar behavior, with a mean duration of the interaction of 221 min or 220 min, respectively. This was markedly longer than for partial contacts which lasted 162 min for dots and 38 min for donuts (Figure 3E). The spreading time of NK cells forming complete synapses, i.e. the duration between the initial contact with the AIS and complete coverage of the print, was significantly longer on donut-compared to dot-shaped AIS (Figure 3F). Thus, the spatial distribution of αCD16 and αLFA-1 influenced the fraction of NK cells forming mature immune synapses by altering the process of NK cell spreading across the AIS. However, for NK cells reaching complete spreading the contact stability was similar for dot- and donut-shaped AIS.

**Figure 3.**
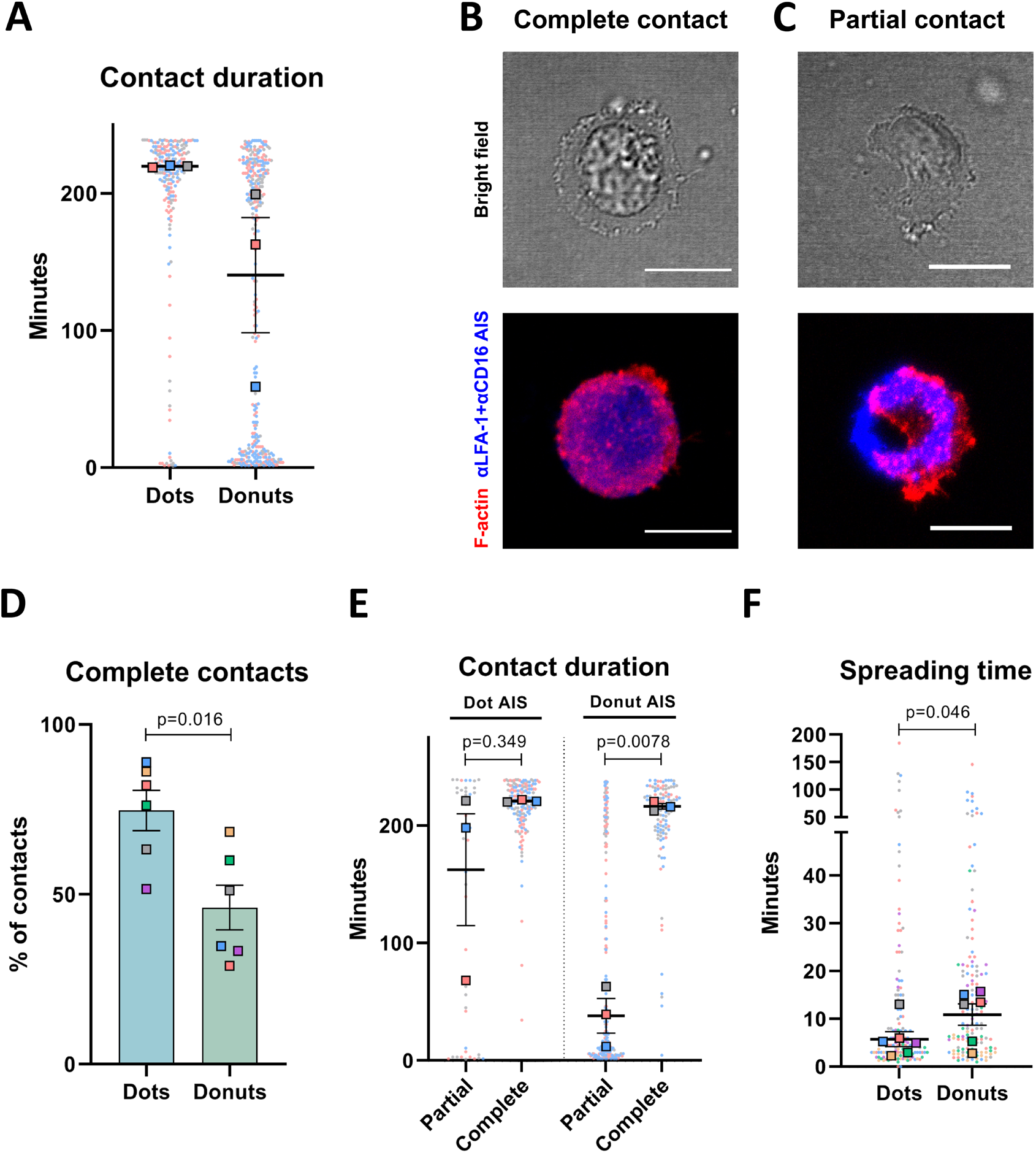
NK cells more commonly build short-lived, partial contacts on donut-shaped AIS compared to dot-shaped αLFA-1 + αCD16 AIS. **(A)** Duration of contacts made by NK cells on dot- or donut-shaped αLFA-1 + αCD16 AIS. Only contacts formed during the first 60 min were included, with a total assay time of 240 min. **(B-C)** Representative images of NK cells in complete (B) or partial (C) contact on donut-shaped αLFA-1 + αCD16 AIS. NK cells were labeled for f-actin (red). **(D)** Proportion of contacts where the NK cell reached complete, symmetric coverage of dot- or donut-shaped αLFA-1 + αCD16 AIS. **(E)** Duration of partial and complete contacts on dot- or donut-shaped αLFA-1 + αCD16 AIS. **(F)** Spreading time of NK cells forming complete contacts on dot- and donut-shaped αLFA-1 + αCD16 AIS, measured as the time between initial contact and reaching complete, symmetrical spreading over the AIS. Data in (A, E) from 3 independent experiments, N=60-90 cells per condition and experiment. Data in (D, F) from 6 independent experiments, N=60-90 cells per condition and experiment. Scale bars: 10 μm.

### NK cell spreading on αLFA-1 + αCD16 AIS correlates with calcium response

Migration arrest upon antigen-recognition of thymocytes has been shown to be associated with transient increases in intracellular calcium (Bhakta et al., 2005). To investigate how NK cell recognition and spreading was governed by calcium signaling, NK cells were loaded with the calcium sensitive dyes Fluo-4 and FuraRed and imaged while interacting with dot- and donut-shaped AIS of αCD16 and αLFA-1. We observed strong calcium fluxes upon synapse formation with AIS and throughout the spreading process, confirming NK cell activation by the AIS (Figure 4A, Supporting Movies 4-5). Most calcium fluxes took the shape of a steep main calcium peak, sometimes followed by a brief drop in fluorescent intensity and one or several secondary peaks, and a slower decay down to a plateau, commonly involving calcium oscillations (Figure 4B). In accordance with previous reports (Leibson et al., 1990), the timing of the onset of the calcium flux was most often associated with morphological changes indicating NK cell commitment to forming a synapse, i.e. cell body polarization and membrane spreading at the contact (Figure 4C). A small fraction of the NK cells did not show a calcium signal upon contact with AIS. These cells almost exclusively formed short-lived partial contacts lacking NK cell commitment, while the vast majority of cells forming committed partial or complete contacts showed a calcium signal (Figure 4D). Comparing NK cells forming partial or complete contacts on either AIS, we observed higher calcium amplitudes for NK cells that went on to spread out over the entire AIS (Figure 4E). Also, the spreading time of NK cells forming complete contacts inversely correlated with the amplitude of the initial calcium peak (Supporting Figure S2 C). We further characterized the decay profile of the calcium fluxes for individual cells by measuring the “sustained fraction”, defined as the fraction of time points where the calcium signaling intensity was higher than half of the peak amplitude, over 20 minutes (i.e. 60 imaging frames) following the initial calcium peak (Supporting Figure S2 D). Although sustained fractions were slightly higher in cells forming complete contacts on either AIS, no correlation was observed between sustained fraction and spreading time across the AIS (Supporting Figure S2 E and F). These results show that calcium signaling is crucial for NK cell engagement, with the amplitude of the initial calcium peak, rather than sustained signaling, dictating the spreading response and formation of a mature immune synapse.

**Figure 4.**
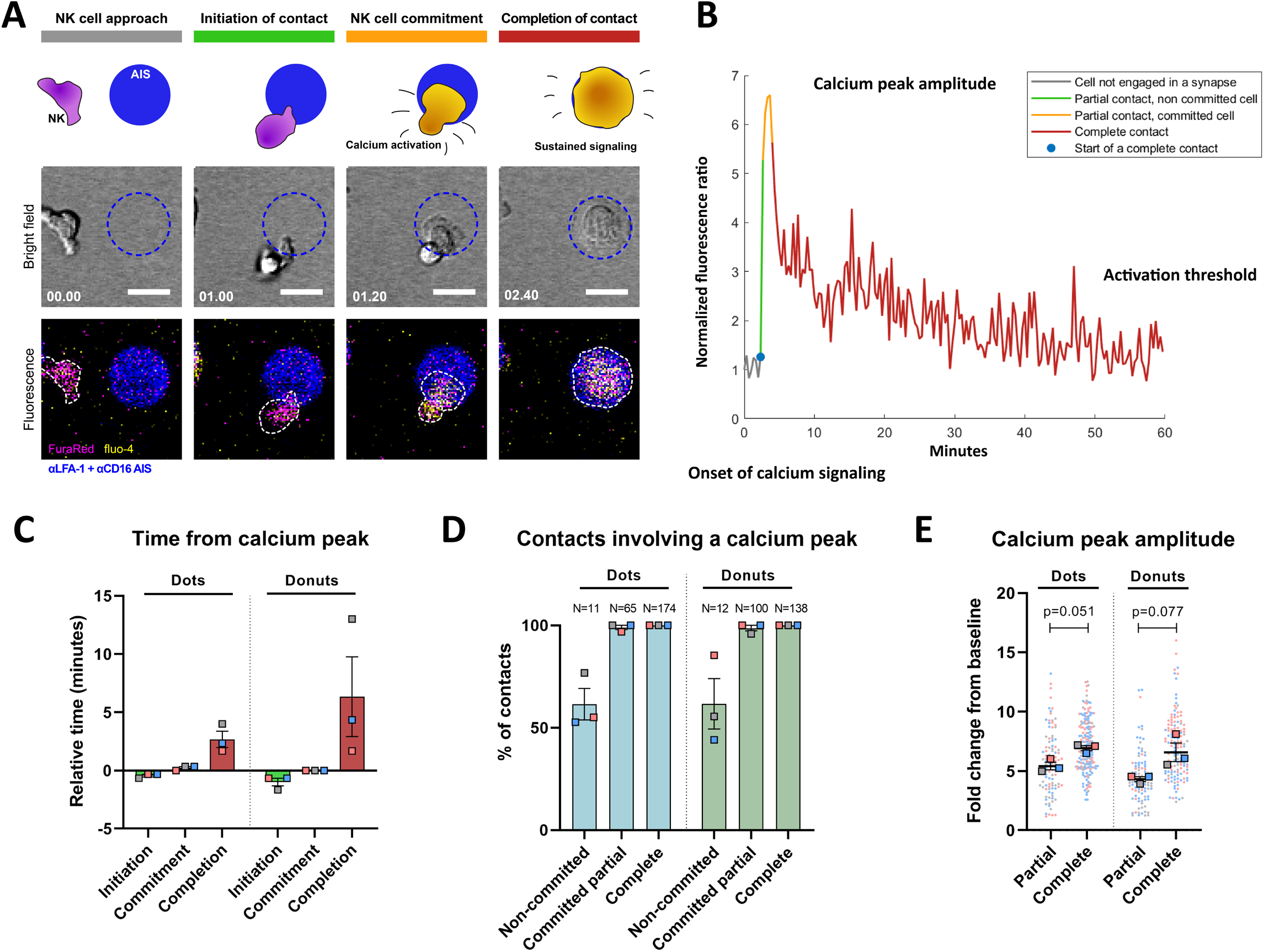
NK cells’ calcium response on αLFA-1+αCD16 AIS coincide with the timing of commitment of the cell to the synapse, and the amplitude of the response correlate with the ability to spread across the AIS. **(A)** Time-lapse sequence of an NK cell building a complete contact on a dot-shaped αLFA-1 + αCD16 AIS. Top row: schematic view of the main stages of interaction as defined by NK morphological changes, approach, initiation of contact, commitment to the synapse, and completion of the contact when the cell evenly covers the AIS. Middle and bottom rows: Bright-field and fluorescence images showing the correlation between morphological changes and calcium activation. The time in minutes elapsed since the left-most image is indicated in white. **(B)** Calcium activity curve of the NK cell during the interaction depicted in (A), corresponding to the normalized ratio between the fluo-4 and FuraRed fluorescent signals. The different stages of the interaction are color-coded as indicated. **(C)** Relative time between the different interaction stages and the onset of calcium activity. Negative values indicate that the interaction stage happened prior to the onset of calcium signaling. **(D)** Proportion of contacts exhibiting calcium activity above the activation threshold, sorted depending on the interaction stage reached: “complete” if symmetric spreading was achieved, “committed partial” if NK cell commitment to the synapse could be observed or “non-committed” otherwise. **(E)** Calcium peak amplitude of NK cells reaching complete or partial contact with either AIS shape, defined as the fold change between the calcium activity at peak and at baseline (prior to contact initiation). Data in (C), (D) and (E) from 3 independent experiments. N=70-100 per condition and experiment. Scale bars: 10 μm.

### NK cells form tight synapses with αCD16 + αLFA-1 AIS despite the local depletion of ligands

Previous studies have described the formation of tight immune synapse with a narrow synaptic cleft spanning up to 35nm (McCann et al., 2003; Stinchcombe et al., 2001). This has also been observed in artificial synapses established on coated surfaces or lipid bilayers (Cartwright et al., 2014). To assess the distance of the synaptic interface formed between the NK cell and the AIS, total internal reflection fluorescence (TIRF) microscopy was used. This technique only excites fluorescence within a very narrow section (≈ 130 nm) closest to the glass substrate and the AIS. NK cells having interacted for 30 min with αLFA-1 + αCD16 AIS were loaded with the membrane dye CellMask Deep Red and imaged by TIRF (Figure 5A and B). Selecting NK cells that had formed complete synapses on dot or donut AIS, we observed sustained membrane fluorescent intensity across both types of prints, showing that the NK plasma membrane laid flat on the entire AIS surface, *i.e.* within 130 nm from the glass surface. For a more detailed assessment we analyzed the radial distribution of fluorescence across the AIS (Figure 5C and D). For donut-shaped AIS, the fluorescence intensity was slightly higher on the ring of antibodies but relatively even in the central region, confirming that the NK cell membrane is close to the entire AIS, despite the lack of specific ligand engagement in the inner region (Figure 5D). Because the TIRF evanescent wave intensity decreases exponentially from the glass surface, intensity variations in TIRF images can be read as differences in distances from the surface for samples with evenly distributed fluorophores (Truskey et al., 1992). The intensity profile on donut-shaped AIS is thus in good accordance with other descriptions of the synaptic cleft. However, exact distance measurements are difficult in our assay because membrane ruffling can also result in locally higher TIRF intensities (Truskey et al., 1992). These results suggest that the complete spreading and formation of a tight synaptic cleft of NK cells over dot- and donut-shaped AIS containing αLFA-1 and αCD16 is not only the mechanical consequence of local ligand engagement. Rather, NK cells integrate the signals from spatially separated ligands and respond by establishing a synapse, where tight contact can be maintained despite the local void of ligands.

**Figure 5.**
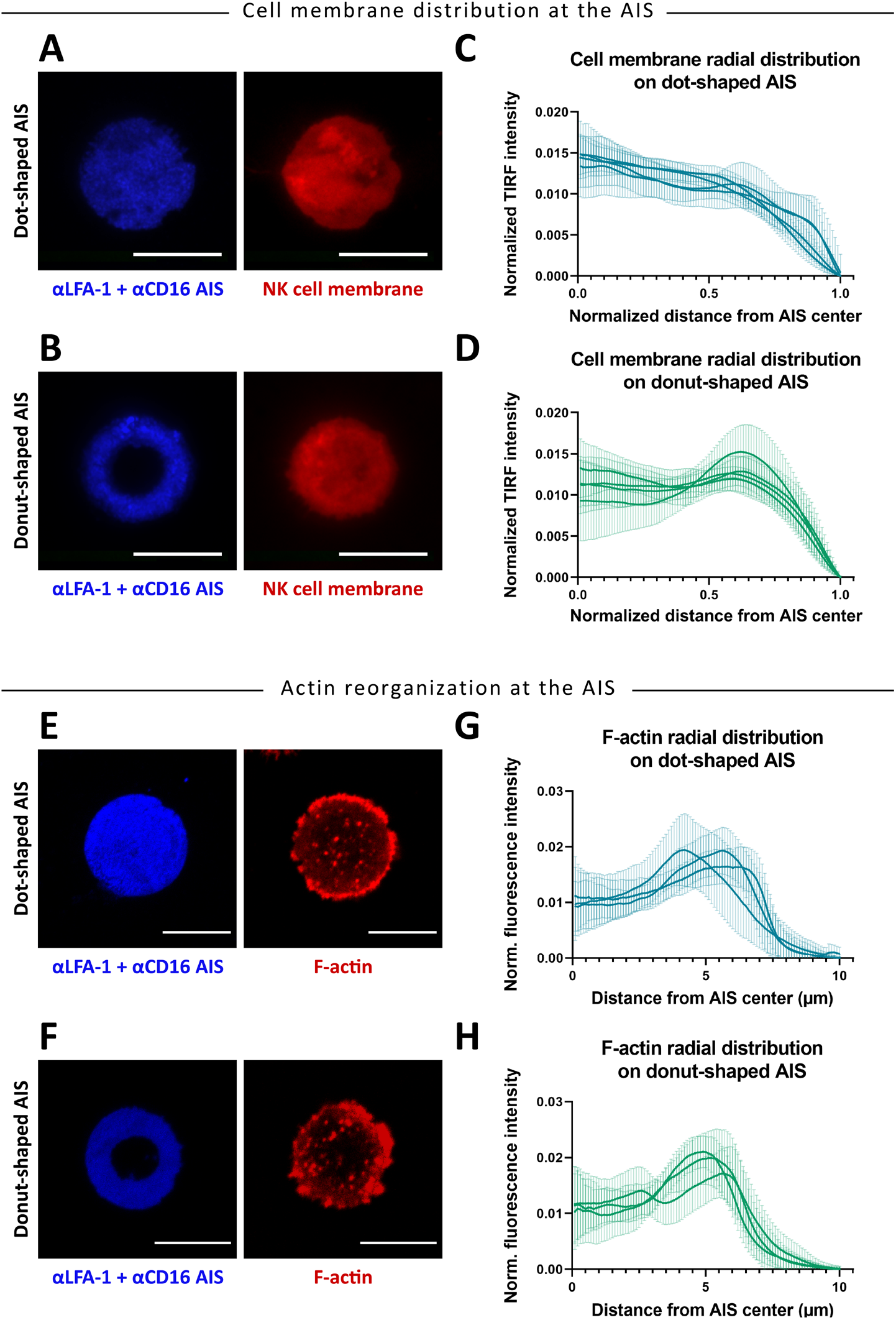
NK cells form a tight contact surrounded by a region enriched in F-actin on both dot- and donut-shaped αLFA-1+αCD16 AIS. **(A, B)** Representative TIRF images of the NK cell membrane during complete contacts on dot-(A) or donut-shaped AIS (B). **(C, D)** Cell membrane radial distribution profile of NK cells on either dot-(C) or donut-shaped AIS (D). Individual curves represent independent experiments, N=16 per condition and experiment. Scale bars: 10μm. **(E, F)** Representative confocal images of F-actin in NK cells forming complete contacts on dot-(E) or donut-shaped AIS (F). **(G, H)** F-actin radial distribution profile of NK cells on either dot-(G) or donut-shaped AIS (H). Individual curves represent independent experiments, N=50-125 per condition and experiment. Scale bars: 10 μm.

### Symmetric spreading of NK cells is coupled to peripheral actin polymerization and radial distribution of PKC-⊝ microclusters

The correlation we observed between symmetric spreading and longer contact duration is in line with previous suggestions that radial symmetry is important for the stability of the immune synapse, in particular *via* actin enrichment in the periphery (Sims et al., 2007; Verkhovsky et al., 1999). Imaging of NK cells fixed and stained for f-actin after 40-minute incubation on αLFA-1 + αCD16 AIS showed that cells that had spread symmetrically had increased actin polymerization at the periphery of the contact (Figure 5E and F).

PKC-⊝ is a principal kinase in T cell signaling and has been shown to redistribute to the immune synapse upon T and NK cell activation (Merino et al., 2012; Monks et al., 1997). Since PKC-⊝ has been shown to break synapse symmetry and promote migration in T cells (Doh & Irvine, 2006; Sims et al., 2007), we investigated the localization of PKC-⊝ in NK cells that had formed complete contacts on αLFA-1 + αCD16 AIS of either shape. In accordance with previous observations in NK cells (Merino et al., 2012), we found that PKC-⊝ organized into microclusters close to the synaptic interface (Supporting Figure S3 A-B). A radial analysis of the fluorescent intensity revealed that PKC-⊝ was principally distributed in an annular structure at the interface between the cell body and lamellopodia, on both dot- and donut-shaped AIS (Supporting Figure S3 C-D). This corresponds to the junction between pSMAC and dSMAC, which also correlates with the distribution of PKC-⊝ in naïve T cells interacting with activating planar bilayers (Sims et al., 2007). The intensity of PKC-⊝ was generally higher around the MTOC as shown by directional analysis of the center of mass of intensity for MTOC and PKC-⊝ (Supporting Figure S3 E-F), which is consistent with a role in positioning of the MTOC at the synapse (Quann et al., 2012). For NK cells forming asymmetric partial contacts with αLFA-1+αCD16 prints, PKC-⊝ was often distributed around the MTOC but could also be found at other locations of the cells (Supporting Figure S3 G-H). Overall the distribution appeared less regular in elongated NK cells compared to NK cells symmetrically spread on prints. Thus, PKC-⊝ assumed a circular distribution with particular weight around the MTOC in NK cells evenly spread across AIS of αCD16 and αLFA-1.

### NK cells rearrange the MTOC and lytic granules by integrating spatially separated signals

Next we set out to investigate how the spatial distribution of ligands influences the polarization of the lytic machinery towards activating AIS. To analyze this, NK cells interacting with dot and donut-shaped αCD16 + αLFA-1 AIS were fixed and stained for microtubules and perforin (Figure 6A). NK cells that had spread symmetrically across the AIS were selected for analysis and the positions of the MTOC and lytic granules relative to the center of the AIS were determined (Figure 6B). In a vast majority of NK cells, the MTOC was polarized (z-direction) towards the AIS (Figure 6C) and slightly laterally displaced (xy-plane) from the center of the AIS (Figure 6D). These observations are consistent with previous reports that the MTOC is not perfectly centered in the cytotoxic immune synapse (Angus & Griffiths, 2013; Stinchcombe et al., 2006). No difference could be observed in lateral positioning of the MTOC between dot and donut AIS (Figure 6D). The MTOC was found within the central, empty region in 74% of complete contacts on donut-shaped AIS (Supporting Figure S4 A-B). Thus, positioning of the MTOC seems to be regulated by integrated signals from the entire AIS rather than by local signaling from bound receptors.

**Figure 6.**
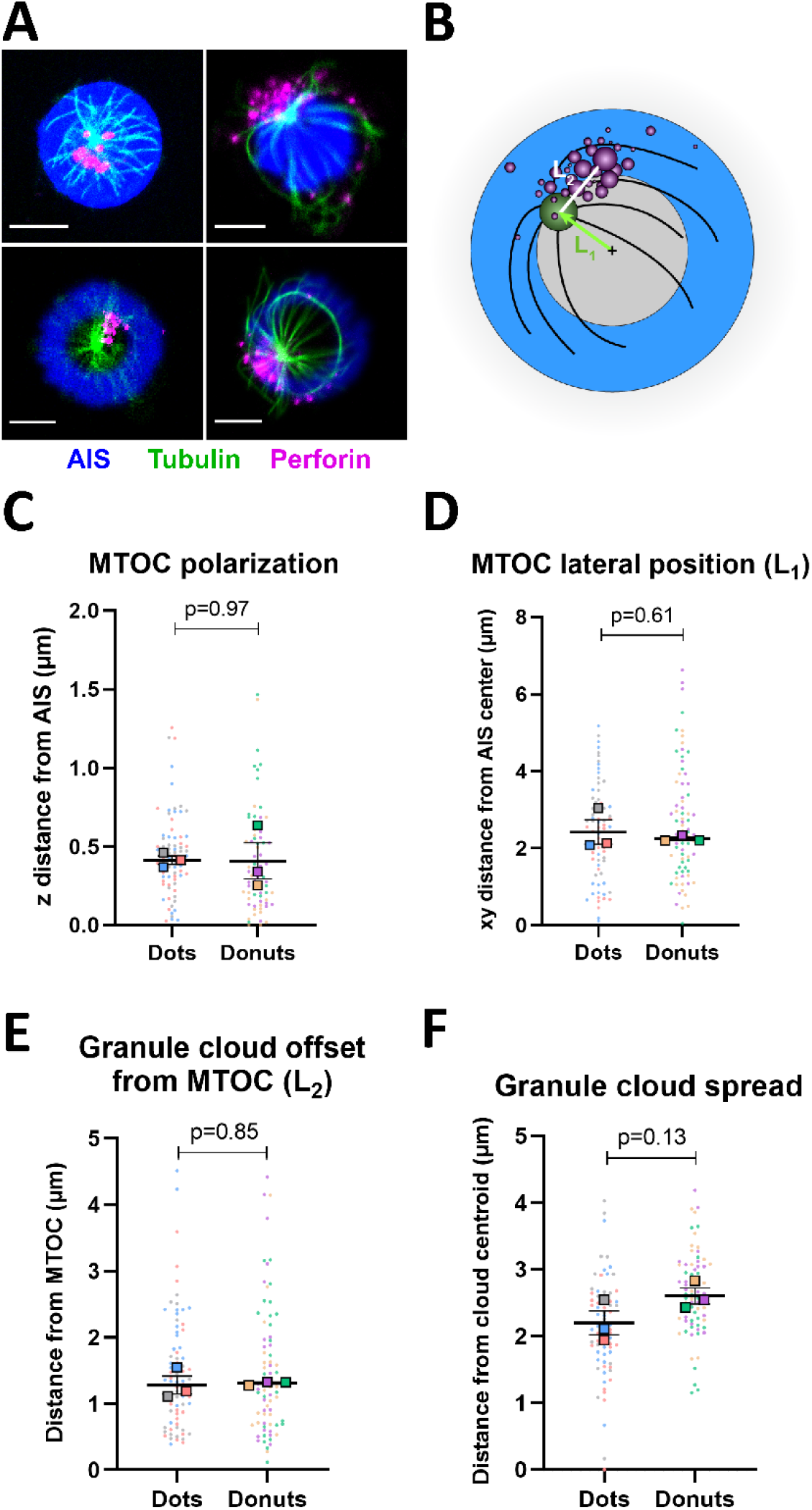
The distribution of αLFA-1 and αCD16 in dot- or donut-shaped AIS has little influence on the organization of the lytic machinery. **(A)** Representative fluorescence images of cytotoxic granules (labelled with anti-perforin. magenta) and microtubules (green) in NK cells building complete contacts on either dot-(top panel) or donut-shaped (bottom panel) αLFA-1 + αCD16 AIS. **(B)** Schematic representation of the parameters describing the spatial organization of granules and MTOC. **(C)** Axial MTOC polarization on dot- and donut-shaped AIS, defined as the distance between the MTOC and the plane of the AIS. **(D)** MTOC lateral position, defined as the x,y distance between the MTOC and the center of the AIS, denoted L1 in (B). **(E)** Granule cloud offset from MTOC, defined as the distance between the MTOC and the centroid of the granule cloud, denoted L2 in (B). **(F)** Granule cloud spread, defined as the average distance between individual granules and the centroid of their cloud. Data from 6 independent experiments, N=25 per condition and experiment. Scale bars: 5 μm.

We next determined the position of the lytic granules with respect to the MTOC. For both AIS shapes, the lytic granules were most often found in a tight cluster with a few satellites. For both dot- and donut-shaped AIS, lytic granules localized close to the MTOC, with the cloud centroid found less than 2 μm away from the MTOC (Figure 6E). The average distance between individual lytic granules and the centroid of the cloud was approximately 2-3 μm (Figure 6F), corresponding well with results found for NK cells activated through interactions with target cells (James et al., 2013). We noticed that the granule cloud was slightly more spread out on donut-compared to dot-shaped AIS (Figure 6F), which could be traced back to a higher amount of granules in the cells (Supporting Figure S4 C-D). The MTOC and granule cloud were however positioned similarly on both AIS shapes, with the granule cloud found most commonly on the outer side of the MTOC relative to the center of the AIS (Supporting Figure S4 E-G). Taken together, these results confirm that NK cells forming complete contacts on αCD16+αLFA-1 AIS organize their lytic machinery in a manner consistent with mature cytotoxic synapses, with the granules tightly packed around the MTOC and close to the interface. The similarity between dot- and donut-shaped AIS shows that this spatial reorganization is independent of receptor engagement in the center of the contact.

### Synapse formation on dot- and donut-shaped αCD16 + αLFA-1 AIS leads to NK cell degranulation targeted at a site of ligand engagement

Having established that both granule convergence and polarization of the lytic machinery happen for complete synapses formed on either AIS shape, we sought to determine whether these synapses could result in degranulation. For this purpose, glass substrates were coated with capture antibodies against perforin (capture αPrf) prior to microcontact printing with either dot- or donut-shaped AIS containing αLFA-1 and αCD16, supplemented with capture αPrf to ensure capture across the entire AIS (Supporting Figure S5). NK cells were fluorescently labelled with Calcein Green and continuously monitored by time-lapse imaging during their interaction with the AIS. The cells were then enzymatically detached from the surface and captured perforin was labelled using biotinylated detection antibodies followed by fluorescently-tagged streptavidin. This allowed us to correlate NK cell dynamics and contact formation with degranulation resulting from the synapse (Figure 7A, Supporting Movies 6-7). We could measure substantial degranulation from NK cells having formed complete contacts on either AIS shape, confirming that the synapses were indeed mature and cytotoxic (Figure 7B). Interestingly, NK cells degranulated less often in complete contacts on donut-shaped AIS, but the amount of perforin captured for individual degranulations was comparable between AIS shapes (Figure 7B and C). Looking at the spatial distribution of degranulation, we found that released perforin formed a tight cluster on both dot- and donut-shaped AIS (Figure 7D). Dividing AIS into concentric regions (Figure 7E), for donut-shaped AIS, the perforin cluster most often localized on the ring with some granules found in the central region devoid of ligands, whereas it localized more towards the center on dot-shaped AIS (Figure 7F and G). This shows that degranulation is targeted towards areas rich in local signaling. These results confirm the maturity of complete synapses on either AIS, but they also indicate a role for the spatial distribution of ligands in regulating the cytotoxic outcome of the synapse.

**Figure 7.**
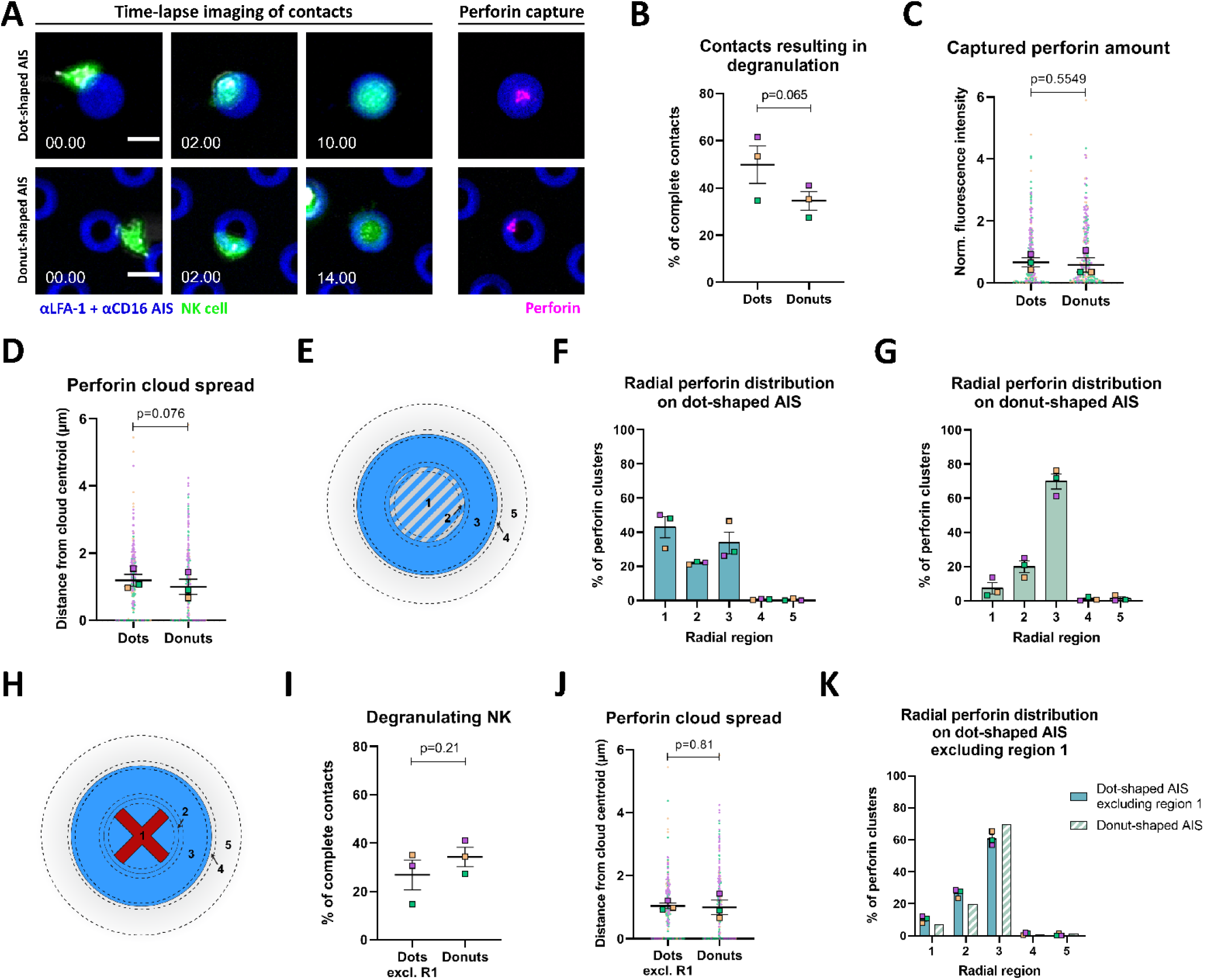
The AIS shape influences the frequency and position of degranulation for NK cells forming complete contacts on αLFA-1+αCD16 AIS. **(A)** Time-lapse sequences and subsequent perforin capture for NK cells building complete contacts on dot-(top panel) or donut-shaped (bottom panel) αLFA-1 + αCD16 AIS. The time in minutes elapsed since the left-most image is indicated in white. **(B)** Proportion of contacts resulting in degranulation, for NK cells having formed complete contacts on αLFA-1 + αCD16 dot- or donut-shaped AIS. **(C)** Perforin amount captured from NK cell forming complete contacts on dot- or donut-shaped AIS, defined as the normalized integrated perforin fluorescent intensity over the AIS. **(D)** Spread of the cloud of captured perforin, defined as the average distance between individual perforin clusters and the centroid of the perforin cloud. **(E)** Schematic representation of the concentric regions used to describe the spatial distribution of captured perforin. 1. 6μm-wide central region of the AIS, corresponding to the void of ligands for donut-shaped AIS. 2. 1μm-wide ring corresponding to the inner border of donut-shaped AIS. 3. Ring corresponding to the main antibody-covered part of the donut-shaped print. 4. 1μm-wide band at the outer border of the AIS. 5. Rim surrounding the AIS, stretching from the edge of region 4 up to 10 μm from the AIS center. **(F-G)** Spatial distribution of individual perforin clusters into the regions defined in (E). Data from 3 independent experiments, N=80 from each. Scale bars in (A): 10 μm. **(H)** To investigate our hypothesis of hindered degranulation in the central void of ligands on donut-shaped AIS, we simulated the effect of degranulation hindrance in region 1 of dot-shaped AIS. Cells having degranulated to an area centered in region 1 were excluded from the analysis and previous parameters were re-calculated. **(I)** Proportion of NK cells degranulating among NK cells forming complete contacts on centrally hindered dot- or donut-shaped AIS. **(J)** Spread of the cloud of captured perforin on centrally hindered dot- or donut-shaped AIS. **(K)** Spatial distribution of individual perforin clusters on centrally hindered dot- and donut-shaped AIS, according to the regions defined in (E).

Since the lateral organization of granules and MTOC is almost identical between cells on both AIS, the difference in the resulting degranulation profile at the surface must result from a difference in granule fusion at the cell surface upon exocytosis. We hypothesized that the position of the lytic machinery is determined according to the integration of signaling from the entire synapse, but that the release of granules is conditioned by local signaling. In donut-shaped AIS, this would result in granules targeted to the central region not being able to fuse and be released, whereas only granules docking close to ligand interactions would be exocytosed and their content captured at the AIS surface. Comparatively, on dot-shaped AIS, practically all granules would be targeted to regions presenting signaling and could thus be released. To test our hypothesis, we sorted degranulating cells on dot-shaped AIS depending on the location of the centroid of their degranulation cloud, here used as a measure for the targeting of granules. We then excluded cells with perforin clouds aimed at the central region of the print (Figure 7H). This hypothetical situation mimics what would happen on donut-shaped AIS if the granules were targeted exactly as on dot-shaped AIS but were only able to be released in the presence of local signaling around the site of docking. In this situation, the number of degranulating cells is strongly reduced, to levels comparable to that on donut AIS (Figure 7I). The radial distribution profile of degranulation on the remaining cells is also almost identical to that observed on donut-shaped AIS, and so is the spread of the degranulation cloud (Figure 7J and K). Thus, these results support that NK cells use the overall shape of the AIS to position the lytic machinery but require local stimuli at the site of degranulation.

## Discussion

Here we have used microcontact printing to pattern antibodies against NK activating receptors in layouts designed to mimic ligand presentation in the immune synapse. By time-lapse imaging we could study the dynamic response of NK cells making contacts with these AIS. In accordance to previous results involving NK cells interacting with antibodies presented on evenly coated surfaces (Culley et al., 2009) or in lipid bilayers (Grakoui et al., 1999; Gross et al., 2010), we observed that ligation of LFA-1 induces a migratory response with NK cells assuming elongated shapes. In this context of spatially separated synapses, this translated in a fraction of NK cells initiating contacts with several AIS simultaneously, sometimes facilitated by formation of thin membrane tethers, or nanotubes, stretching between distal AIS and the NK cell body. Interestingly, NK cells commonly stayed in contact with a single AIS, while actively seeking for subsequent contacts with the rest of the cell body, revealing a seemingly contradictory consequence of LFA-1 signaling where both tethering and motility are supported in parallel. In contrast, combined ligation of LFA-1 and CD16 led to reduced NK cell migration often followed by spreading across single AIS into symmetrical, stable contacts. This confirms the role of CD16 engagement in inducing NK cell commitment to the synapse, providing a stop signal which balances the motility signals from LFA-1.

We further investigated the importance of local signaling in the immune synapse formation and outcome. For this purpose, we compared the effect of αCD16 + αLFA-1 signaling in synapses with central ligand engagement (dot-shaped AIS) and in synapses with a central depletion of ligands (donut-shaped AIS). We observed that altering the distribution of ligands affected the stability of contacts, as NK cells generally formed shorter contacts on donut-shaped AIS. This could be explained by a higher proportion of partial contacts on donut-shaped AIS, where the NK cells were unable to continue to spread across the AIS and thus to cover the central void of ligands. Live cell calcium imaging showed that a fraction of NK cells with high activation response could spread across donut-shaped AIS to establish complete contacts. The difficulty for NK cells to reach a radially symmetrical configuration on donut-shaped AIS without spreading over the entire AIS often resulted in continued motion and eventually detachment, whereas spreading over dot-shaped AIS led to maintained symmetry and thus remarkable stability (Culley et al., 2009; Verkhovsky et al., 1999). Here we used a static model system of immobilized antibodies which is obviously very different from NK cell spreading across a target cell with a fluid membrane. Nevertheless, our results suggest that not only the nature of ligands but also their distribution across the target cell membrane could influence the outcome of NK cells surveillance by promoting or inhibiting NK cell spreading, possibly by altering the symmetry of the synapse which could result in resumed migration (Mayya et al., 2018; Verkhovsky et al., 1999).

We then focused on the fraction of NK cells that went on to form complete contacts over the AIS. We found that their morphology, contact stability, calcium signaling profile, plasma membrane organization at the interface, and positioning of the MTOC and perforin-containing granules were largely similar between dot- and donut-shaped AIS. NK cells forming complete contacts showed robust calcium signals upon commitment to the contact with the AIS and once spread out had a symmetric, round morphology with actin accumulation at the cell periphery. The synapses proved to be very stable with contacts often lasting for several hours. TIRF imaging confirmed that the NK cell plasma membrane was in close proximity with the whole AIS surface with some local variation in intensity, suggesting either tighter contact or increased accumulation of membrane in the center for NK cells on dots and on top of the antibody ring for NK cells on donut-shaped AIS. In NK cells that had been fixed 1 hour after seeding, the MTOC surrounded by a tight cluster of perforin-granules was commonly found within 2 μm from the AIS surface on both dot- and donut-shaped AIS, confirming cytotoxic commitment of the NK cells (Bryceson et al., 2005; Chen et al., 2006; Sancho et al., 2000). Also, the granule cloud and MTOC showed similar lateral organization on both AIS shapes, with the MTOC placed off-center, approximately 2.5 μm away from the center of the AIS. Thus, on donut-shaped AIS, the MTOC was most often found over a region devoid of ligands.

Taken together, the similarities observed between dot- and donut-shaped AIS indicate that once a complete contact is formed over the AIS, the assembly of the immune synapse is independent of the difference between dot- and donut-shaped prints. NK cells thus sense the overall circular shape of the print rather than the central part. We believe that the circular shapes of the AIS promoted maintained symmetric spreading by facilitating a directional balance between the centripetal contractile forces (Culley et al., 2009; Döbereiner et al., 2006; Sims et al., 2007; Verkhovsky et al., 1999; reviewed in Dustin, 2007). We investigated the distribution of PKC-⊝, a principal kinase with a suggested role in synapse symmetry in T cells (Monks et al., 1997; Sims et al., 2007; Zanin-Zhorov et al., 2011; reviewed in Dustin, 2009). In NK cells having spread evenly on either AIS, the immune synapse stability could be supported by the symmetric and radial distribution of PKC-⊝ at the junction between p- and dSMACs, a distribution that was not found in NK cells that only partially covered an AIS or migrated on the glass between AIS. This observation fits with the general picture that the immune synapse, where the effector cell spreads symmetrically across the target cell and activating receptors and ligands are accumulated into circularly shaped clusters, is a structure that promotes stability, sustained signaling and preparation for effector functions.

Next we used a combination of live-cell time-lapse imaging and antibody capture at the glass surface to associate NK cell interaction with the AIS with the resulting degranulation. We measured substantial degranulation from NK cells forming complete contacts on both dot- and donut-shaped AIS, confirming that NK cell interaction with AIS could mimic NK-target cytotoxic synapses. Fewer cells degranulated in complete contacts on donut-shaped AIS compared to dots, which suggests a link between ligand distribution and cytotoxic outcome of the synapse. This concept was recently proposed for breast cancer cells where the induction of an “actin response” in reaction to NK cell recognition resulted in reduced cytotoxicity, possibly *via* modulation of the local concentration of NK cell ligands (Absi et al., 2018).

Analyzing the radial distribution of captured perforin on the AIS showed that a population of cells degranulating towards the center of dot-shaped AIS was absent on donut-shaped AIS. In fixed NK cells, the lateral distribution of granules and the positioning of the MTOC were almost identical between cells on both AIS, suggesting that the lytic machinery was being targeted to the same region independently of the central composition of the AIS. We hypothesized that the difference in the resulting degranulation profile at the surface could then result from differences in the subsequent steps leading to exocytosis, and that this would be dependent on local signaling. We tested this by excluding NK cells where the centroid of the degranulated perforin cloud was found in the center of the dot-shaped AIS from the analysis. This brought down the hypothetical fraction of degranulating NK cells down to a similar level as that observed on donuts, and the radial distribution of perforin to a profile similar to that measured on donuts. Thus, this supports the idea that local signaling is required for the final steps leading to CD16 signaling-induced degranulation, in line with the observation that sites of granule release are commonly found in proximity with CD16 microclusters (Steblyanko et al., 2015). The lack of local signaling in the predetermined area geared for degranulation would then result in impaired exocytosis, as observed in the central region on donut-shaped AIS.

## Conclusion

In this study we investigated how NK cells interact with static protein prints sized as typical NK cell immune synapses shaped as dots or donuts. The main conclusions from the study are: 1. Ligand distribution affects the NK cells’ ability to spread across the AIS, indicating that target cells could play an active role in inhibiting or promoting establishment of an immune synapse. 2. For cells establishing a complete contact, stability and assembly of the lytic machinery are regulated by the overall shape of the print, showing that NK cells can sense large-scale patterns and integrate spatially separated signals. 3. Local activating signals are important for the NK cells to proceed with degranulation. Thus, mechanisms that disrupt ligand assemblies in designated areas of the synapse could be a way for target cells to avoid attack. Therefore, the regulation of ligand dynamics in target cells during immune synapse formation deserves further studies.

## Materials and Methods

### Isolation and culture of NK cells

Human NK cells were isolated from blood from anonymous healthy donors according to local ethics regulations, following either of the following protocols. For experiments involving fixed NK cells on AIS, peripheral blood mononuclear cells (PBMCs) were separated from buffy coats by density gradient centrifugation (Ficoll Paque, GE Healthcare). NK cells were then isolated from PBMCs by negative selection according to manufacturer’s instructions (Miltenyi Biotec). For control of NK cell purity, cells were stained with monoclonal antibodies for CD56-PE (BioLegend, clone MEM-188) and CD3-FITC (BioLegend, clone OKT3) and analyzed with a FACS Caliber cytometer. For all other experiments, NK cells were directly isolated from buffy coats using negative magnetic selection according to manufacturer’s instructions (Stem Cell Technologies). NK cells were then stained using the following antibodies: CD56-PE (BioLegend, clone HCD56), CD3-BV421 (BioLegend, clone UCHT1, or BD Horizon, clone SK7), CD16-APC (BD Pharmingen, clone 3G8), NKp46-PE-Cy7 (BD Pharmingen, clone 9E2/NKp46) and the viability dye BV510 (BD Biosciences), before purity analysis on a FACS Canto II flow cytometer (Becton Dickinson).

NK cells isolated using either protocol were maintained in RPMI cell culture medium containing 10% human AB+ serum, 1% penicillin-streptomycin, 2 mM L-glutamine, 1 mM sodium pyruvate, and 1× non-essential amino acids (all from Sigma Aldrich). NK populations contained over 95% CD56+ CD3-cells and were used within 48 hours after isolation.

### Microcontact printing

Stamp masters for printing were produced by etching patterns in silicon using a previously described process (Frisk et al., 2011). Stamps of poly(dimethylsiloxane) (PDMS; Sylgard 184, Dow Corning) were produced by casting prepolymer solution in silicon masters and curing at 65°C for at least 6 hours. To improve the uptake of the loading solution by the hydrophobic PDMS surface, stamps were first washed with ethanol, then washed and degassed twice in PBS. Stamps were then incubated for 1 hour with either 1) 10 μg/mL antibodies against LFA-1 (αLFA-1; BioLegend, clone HI111) or 2) 10 μg/mL αLFA-1 + 10 μg/mL antibodies against CD16 (αCD16; BioLegend, clone 3G8). All loading solutions were mixed in PBS and supplemented with 10 μg/mL bovine serum albumin (BSA) conjugated to Alexa Fluor 555 (BSA-AF555; Thermo Fisher Scientific) for visualization. The stamps were then briefly washed with PBS and MilliQ, dried and placed on either a 35-mm glass-bottom dish pre-coated with poly-D-lysine (MatTek) or an 8-chamber glass slide (ibidi) coated with poly-L-lysine (Thermo Fisher Scientific), with the structured surface facing the glass. Two types of prints were used, either dot- or donut-shaped. The outer diameter of dot-shaped stamps was between 8 and 15 μm, while the inner and outer diameters of donut stamps were 6-7 and 14-15 μm, respectively. For experiments involving AIS of both shapes, outer AIS diameters were matched.

### Live cell migration and calcium imaging

For migration analysis, NK cells were washed twice in PBS then incubated in 1 mL RPMI with 1 μM Calcein Green (Thermo Fisher Scientific) for 20 minutes at 37°C, 5% CO_2_. For calcium imaging, cells were washed twice with HBSS (Invitrogen) before staining with 3 μM fluo-4 (Invitrogen) and 4 μM Fura Red (Thermo Fisher Scientific) for 30 minutes in RPMI. After washing in either PBS or HBSS, NK cells were left to rest for another 30 minutes before imaging. The cells were seeded at 2-3 × 10^5^ cells/mL onto micropatterned glass in complete RPMI, supplemented with 10 mM Hepes (Sigma Aldrich) for migration studies. Imaging was conducted using a 20× Plan-Apochromat objective on a LSM 880 confocal microscope (Carl Zeiss AG) with an incubation chamber set to 37°C, 5% CO_2_, capturing one frame every 45-60 s for 4 hours (migration) or every 20 s for 90 minutes (calcium signaling).

NK cell segmentation and single cell tracking were performed using the “Pixel classification” and “Object tracking with learning” modules in Ilastik (Berg et al., 2019). For all duration measurements (contact duration, spreading time), only contacts formed within the first hour of the four-hour assay were included to avoid measurement artefacts due to cells landing at the surface later in the assay. Different modes of NK cell migration were defined according to the mean square displacement (MSD) along the migration track as previously described (Khorshidi et al., 2011). Briefly, the MSD was evaluated using a sliding window of 20 minutes. Transient Migration Arrest Periods (TMAPs) were identified by comparing the mean diffusion coefficient along the curve with the random diffusion coefficient estimated for a spherical particle of size comparable to a cell. Directed migration was detected by fitting the MSD to *t*^*α*^. *α* = *1* corresponds to a situation of random Brownian motion, and directed migration was characterized as *α* > *1.5* for at least 10 consecutive frames.

Calcium signaling curves were obtained by dividing the mean fluo-4 (fluorescent in its calcium bound form) fluorescence intensity by the mean FuraRed (fluorescent in its calcium free form) fluorescence intensity in individual tracked NK cells, at each time point. The resulting curves, here denoted as calcium intensity curves, were then normalized to their value at the first time point, prior to synapse formation. To define an activation threshold for all cells, potential calcium intensity peaks were identified by comparing the calcium intensity at each time point to the average of all previous time points (fold change). Receiver operating characteristic (ROC) curves were then calculated using NK cells that did not interact with any AIS (true negatives) and a representative sample of NK cells exhibiting a clear calcium intensity peak (true positives). The threshold was chosen to maximize the probability of detection (true positive rate) while minimizing the probability of false alarm (false positive rate) (Salles et al., 2013), leading to a threshold value of 2.63.

Contact times were manually measured in ImageJ (NIH), while individual migration tracks, mean square displacement and calcium signaling curves were analyzed in Matlab (MathWorks).

### High resolution confocal imaging of fixed NK cells on AIS

NK cells were seeded on the micropatterned glass bottom of 35-mm dishes (MatTek) at 5 × 10^5^ cells/ml and were incubated for 40 minutes at 37°C and 5% CO_2_ in RPMI medium containing 1% human serum. Cells were fixed and permeabilized for 20 minutes in Fix/Perm solution (BD Biosciences) and washed using Perm/Wash buffer (BD Biosciences), followed by a blocking step with PBS supplemented with 5% goat serum for 60 minutes. For respective experiments, the cells were then stained with phalloidin conjugated to Abberior STAR 635 (Abberior), antibodies for α-tubulin conjugated to Alexa Fluor 488 (Millipore, clone DM1A) and for perforin conjugated to Pacific Blue (BioLegend, clone dG9). For PKC-⊝ labelling, the cells were first stained with a primary rabbit antibody (Santa Cruz, polyclonal) followed by a goat anti-rabbit secondary antibody conjugated to Alexa Fluor 405 (Thermo Fisher Scientific). Cells were washed and images were acquired using a 63×/1.40 Plan-Apochromat oil immersion objective on a LSM 880 confocal microscope (Carl Zeiss AG).

The contact area of NK cells on AIS was obtained by thresholding the phalloidin fluorescence intensity in the z-plane of the print, followed by topologically closing and filling holes in the obtained mask. Similarly, NK cell roundness was measured on thresholded maximum intensity projections of the phalloidin channel. To determine the position of the MTOC and lytic granules relative to the AIS center on complete contacts, single NK cells with roundness over 0.8 covering at least 90% of the AIS were selected. The position of the MTOC was manually set as the convergence point of microtubules, often corresponding to the α-tubulin cluster of highest intensity in the cell. Lytic granules were detected by segmenting clusters of perforin, and the position of each was defined as the fluorescence intensity-weighted center of mass. The granule cluster centroid was defined as the cumulated intensity-weighted center of mass of all individual granules in the cell. The average distances between individual granules and other objects were always weighted to granule cumulated intensity, thus compensating for segmenting effects (Mentlik et al., 2009). The relative position of the MTOC and the granule cluster was analyzed by measuring the angle between two vectors, both starting from the AIS center, but one drawn to the MTOC and the other to the granule cluster centroid.

NK cell contact area on prints and roundness were measured using ImageJ (NIH). Segmentation of granules was performed in Volocity (PerkinElmer) and distance calculations were performed in Matlab (Mathworks). Radial fluorescence intensity profiles of F-actin and PKC-⊝ were generated from single slice images (F-actin) or maximum intensity projection images (PKC-⊝) using the Radial Profile plugin in ImageJ (NIH).

### TIRF imaging of fixed NK cells on AIS

NK cells were seeded on the micropatterned glass bottom of 35-mm dishes (MatTek) at 5 × 10^5^ cells/ml and left to interact for 30 minutes. NK cell outer membranes were then stained with CellMask Deep Red (Thermo Fisher Scientific) at 10 μg/mL for 10 minutes, washed with PBS before 10-minute fixation with Cytofix (BD Biosciences) containing 4% paraformaldehyde. The samples were then blocked using PBS with 5% w/v BSA and finally mounted for immediate imaging. Images were acquired using an Elyra S1 microscope equipped with an α Plan-Apochromat 100×/1.46 oil immersion objective (Carl Zeiss AG). NK cells having spread symmetrically over the entire AIS were selected by widefield imaging before switching to TIRF mode where the imaging focus plane was set according to the AIS.

To assess if NK cells’ plasma membrane were in close proximity to the surface across the AIS, radial fluorescence intensity profiles of the NK cell membrane over the prints were generated using the Radial Profile plugin in ImageJ (NIH). The resulting curves were normalized in radius (0 to 1) and in intensity (integrated fluorescence intensity set to 1) to account for individual staining differences.

### Perforin capture on AIS

Capture surfaces were prepared by successively coating 8-chamber glass slides (ibidi) with poly-L-lysine (Thermo Fisher Scientific) and 5 μg/mL anti-perforin capture antibody mix (capture αPrf; Mabtech, combination of clones Pf-80/164). The resulting surfaces were micropatterned as described before with 10 μg/mL αLFA-1 + 10 μg/mL αCD16 + 5 μg/mL capture αPrf + 10 μg/mL BSA-AF555. NK cells labelled with 1 μM Calcein Green were added to the chambers at 2-3 × 10^5^ cells/mL. During their interaction with AIS, cells were imaged every 2-5 min using a 20× Plan-Apochromat objective on a CellObserver 7 widefield microscope (Carl Zeiss AG). After 60 min, the cells were detached using the Accumax detachment solution (Merck Millipore) at 37°C, 5% CO_2_ for 15 minutes. Cells were washed away with PBS, and complete cell removal was confirmed by visual inspection using a phase contrast microscope (Carl Zeiss AG). The slides were then washed twice with Elisa buffer (PBS + 0.05% Tween-20 + 0.1% w/v BSA). The detection antibody solution composed of PBS + 5 μg/mL anti-perforin conjugated with biotin (clone Pf-344, Mabtech) was added to the slide chambers. After washing with buffer, captured perforin was revealed by labelling with Streptavidin conjugated with Alexa Fluor 647 (BioLegend) at 1 μg/mL in PBS, and the slides were washed again with Elisa buffer solution. The surfaces were then taken back to the microscope, and the slides were re-aligned so that the same regions of interest were used for degranulation measurements as for migration.

The images were analyzed using ImageJ (NIH). Briefly, time-lapse and perforin detection images were combined and spatially aligned to account for stage drift and imprecision when taking the slide off and on the microscope. On AIS where a single NK cell had established a complete contact, degranulation distribution was measured in a 20 μm-wide circular region centered on the print.

### Statistical analysis and data plotting

Following recently published suggestions (Lord et al., 2020), we chose to only compare summary measures (here set to the median of all individual measurements) between independent experiments as opposed to comparing pooled data from the independent experiments. The plots reflect this, as the large colored squares indicate the median measurement value for each independent experiment, and the cluster plots in the background (where applicable) show the corresponding individual measurements. Paired experiments are color-coded. The thicker black line (or bar height where applicable) indicates the mean of summary measures from independent experiments, and the error bars represent the standard error of mean of summary measures. All p-values were calculated using (paired where applicable, unpaired otherwise) Student’s t-test between the summary measures of indicated groups. P-values were calculated and plots were produced using Prism (GraphPad) or Matlab (Mathworks).

## Supporting information

Supporting Movie 5 - Calcium activity on donut-shaped AIS

Supporting Movie 6 - Capture of degranulated perforin on dot-shaped AIS

Supporting Movie 7 - Capture of degranulated perforin on donut-shaped AIS

Supporting Movie 1 - Oscillations between AIS

Supporting Movie 2 - Partial contact

Supporting Movie 3 - Complete contact

Supporting Movie 4 - Calcium activity on dot-shaped AIS

## Acknowledgements

We thank the Önfelt lab and Daniel Davis for discussions and feedback on the manuscript and Daniel Jans for introduction to TIRF experiments. We thank the Swedish Foundation for Strategic Research, The Swedish Cancer Foundation and The Swedish Childhood Cancer foundation for financial support.

