## Supporting Figures 1-5 for "NK cells integrate signals over large areas when building immune synapses but require local stimuli for degranulation"

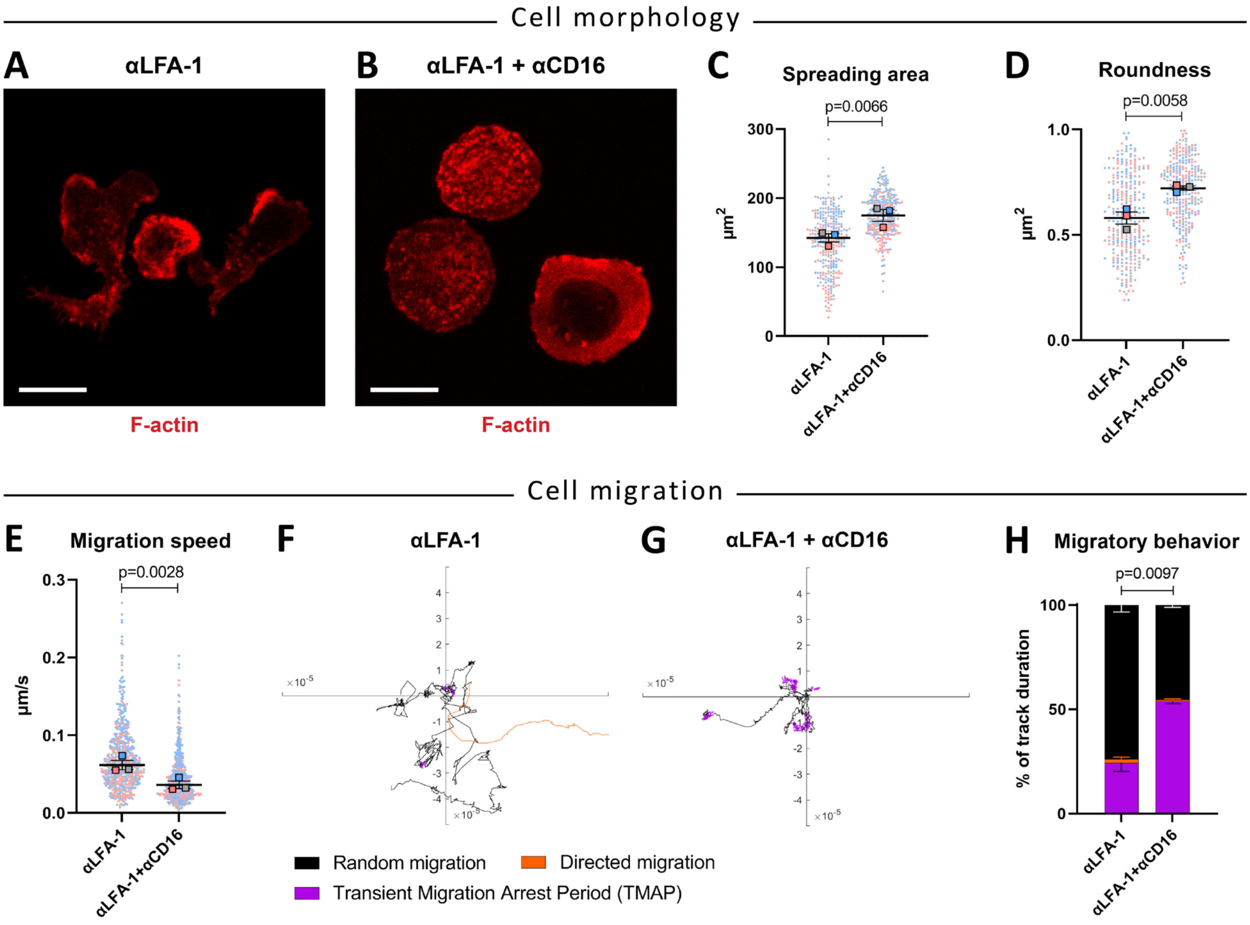


**Supporting Figure S1. Interaction of NK cells with surfaces evenly coated with αLFA-1 results in a migratory behavior, whereas αLFA-1 and αCD16 combined lead to a stop signal and cell spreading.** **(A-B)** Representative fluorescence images of NK cells interacting with glass surfaces evenly coated with αLFA-1 (A) or αLFA-1 + αCD16 (B). **(C)** Spreading area of NK cells interacting with either coated surface. **(D)** Roundness of NK cells interacting with either coated surface. Data in (C, D) are from 3 independent experiments, N=100 per condition and experiment. **(E)** NK cell migration speed on either coated surface. **(F-G)** Example NK migration tracks color-coded for migratory behavior. **(H)** Average percent of time spent in the different modes of migration on either surface. The P-value was calculated by comparing the TMAP fraction between groups. Data in (E, H) are from 3 independent experiments, N=200 per condition and experiment.

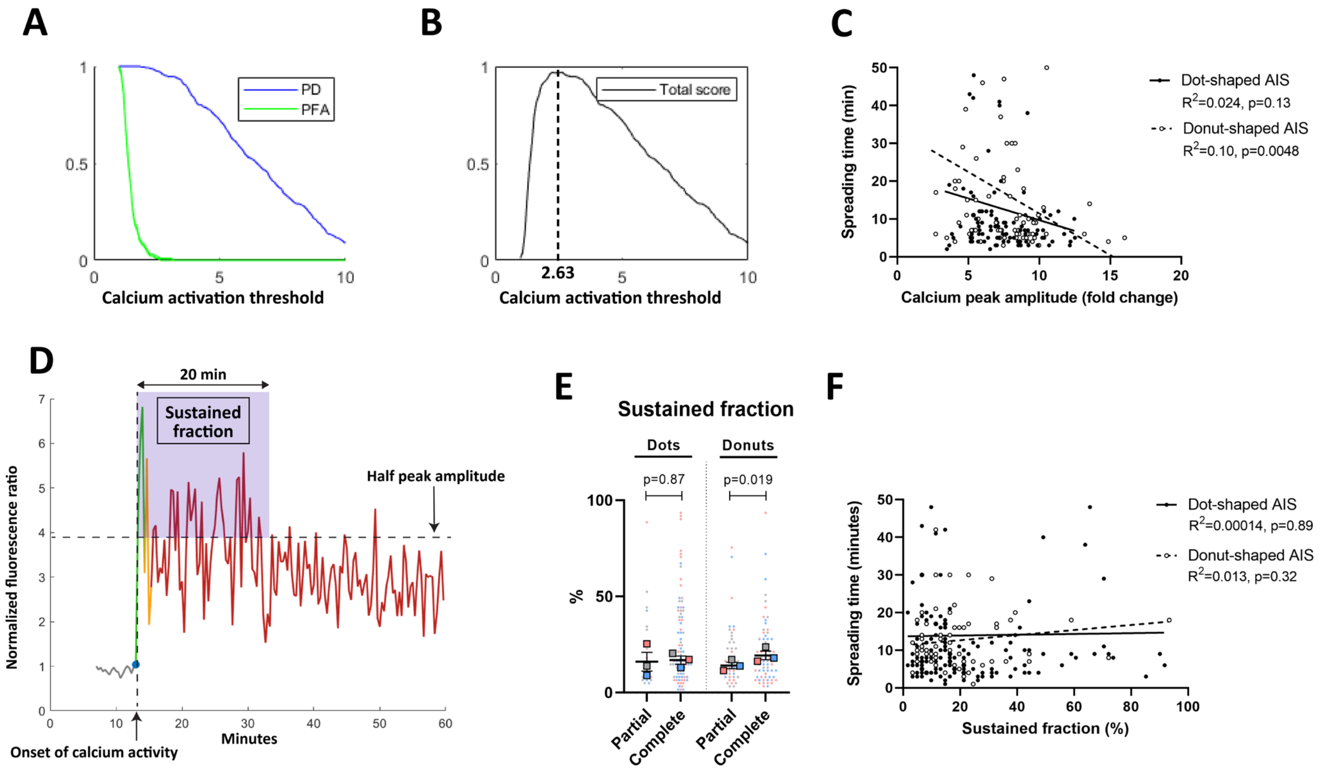


**Supporting Figure S2. NK cell spreading dynamics correlate with the amplitude of calcium activation rather than its decay profile. (A-B)** The calcium activation threshold was determined using receiver operating characteristic (ROC) curves. Using a sample of calcium activity curves from cells that did not interact with any AIS (true negatives) and from cells exhibiting a clear calcium intensity peak (true positives), the probability of detection (PD, true positive rate) and the probability of false alarm (PFA, false positive rate) were calculated (A). The threshold was chosen to maximize the total score, defined as the product of PD and PFA for all tested thresholds, leading to a final activation threshold value of 2.63 (B). **(C)** Correlation between spreading time across the AIS and the calcium peak amplitude measured, for NK cells having formed a complete contact on dot- or donut-shaped αLFA-1 + αCD16 AIS. **(D)** We defined the sustained fraction as the percent of time where the NK cell showed higher calcium activity than a certain threshold, set as half the peak calcium amplitude, over 20 minutes following the onset of calcium signaling. **(E)** Sustained fraction measured for NK cells interacting at least 20 minutes on dot- or donut-shaped AIS. **(F)** Correlation between spreading time across the AIS and the sustained fraction, for NK cells having formed a complete contact on dot- or donut-shaped αLFA-1 + αCD16 AIS.

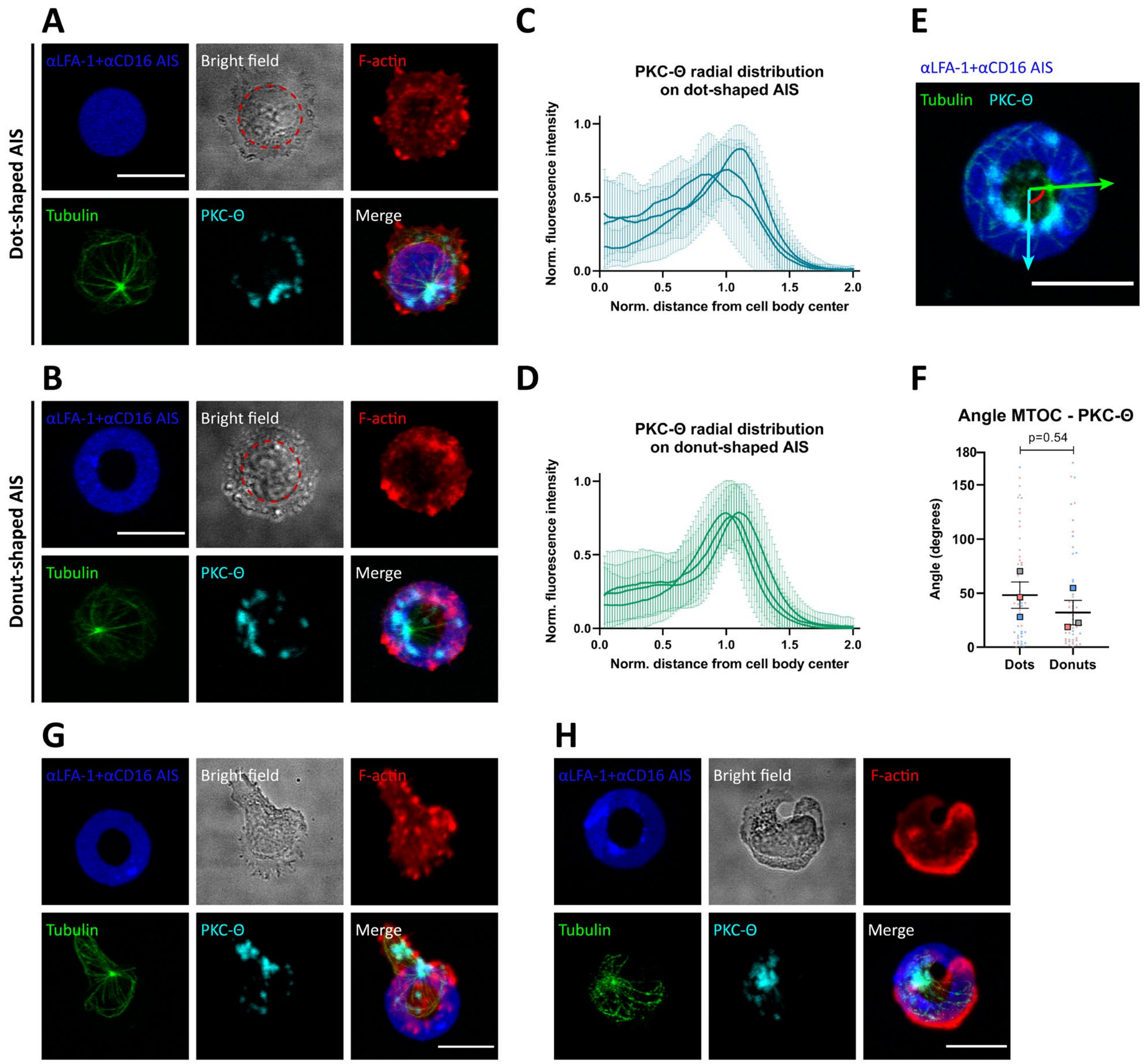


**Supporting Figure S3. PKC-Θ organizes in a ring structure at the border between the cell body and lamellopodia in NK cells forming complete contacts on dot- or donut-shaped αLFA-1 + αCD16 AIS.** NK cells interacting with αLFA-1 + αCD16 AIS in dot or donut shape were fixed and labelled for F-actin (red), tubulin (green) and PKC-Θ (cyan). **(A, B)** Representative fluorescence images of NK cells forming complete contacts over dot- (A) and donut-shaped (B) AIS. **(C, D)** Radial distribution of PKC-Θ in NK cells forming complete contacts on dot- (C) or donut-shaped AIS (D). A distance of 1.0 from the cell body center corresponds to the edge of the cell body, illustrated by a dashed red ellipse in panels (A, B). Individual curves in (C, D) represent independent experiments with average values from N=26-60 cells. **(E)** On NK cells stained for tubulin and PKC-Θ, the angle (red) between two vectors was measured, both stemming from the center of the AIS, one directed towards the MTOC (green) and one towards the centroid of all PKC-Θ clusters in the cell (cyan). **(F)** Angle between the MTOC and the centroid of PKC-Θ clusters relative to the center of the AIS. **(G-H)** Representative fluorescence images of NK cells forming partial contacts on donut-shaped αLFA-1 + αCD16 AIS. Scale bars: 10µm.

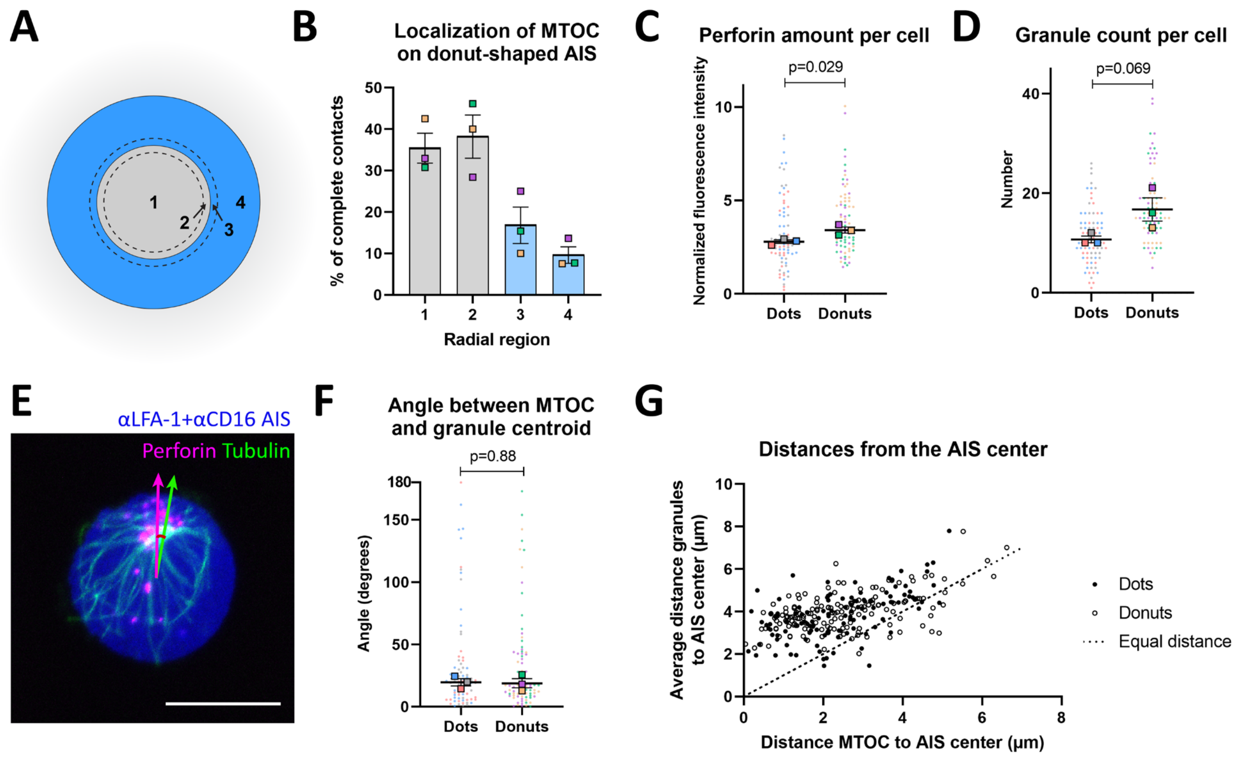


**Supporting Figure S4. The cloud of perforin granules is most often found on the outside of the MTOC relative to the AIS center, in NK cells forming complete contacts with αLFA-1+αCD16 AIS.** NK cells having formed complete contacts with αLFA-1 + αCD16 AIS of either shape were fixed and stained for F-actin, tubulin and perforin, and the respective positions of the MTOC and the centroid of the granule cloud were determined. **(A)** Donut-shaped AIS were divided into concentric regions to analyze the radial positioning of the MTOC. **(B)** Localization of the MTOC in the concentric regions defined in (A). **(C)** Total perforin amount measured per NK cell, defined as the cumulated perforin fluorescent intensity over the entire cell, normalized to the standard deviation of perforin intensity measurements in each independent experiment. **(D)** Granule count per cell, defined as the number of segmented perforin objects per cell. The lower perforin amount and granule count in NK cells on dot-shaped AIS could be a result of more frequent degranulation. **(E)** On NK cells stained for tubulin and perforin, the angle (red) between two vectors was measured, both stemming from the center of the AIS, one directed towards the MTOC (green) and one towards the centroid of perforin granules in the cell (magenta). **(F)** Angle between the MTOC and the centroid of perforin granules relative to the center of the AIS. **(G)** Comparison between the average distance between granules and the center of the AIS, and the distance between the MTOC and the center of the AIS. The dashed line represents an equal distance from the center of the AIS. Data from 3 independent experiments per condition, N=25 per experiment. Scale bar: 10µm.

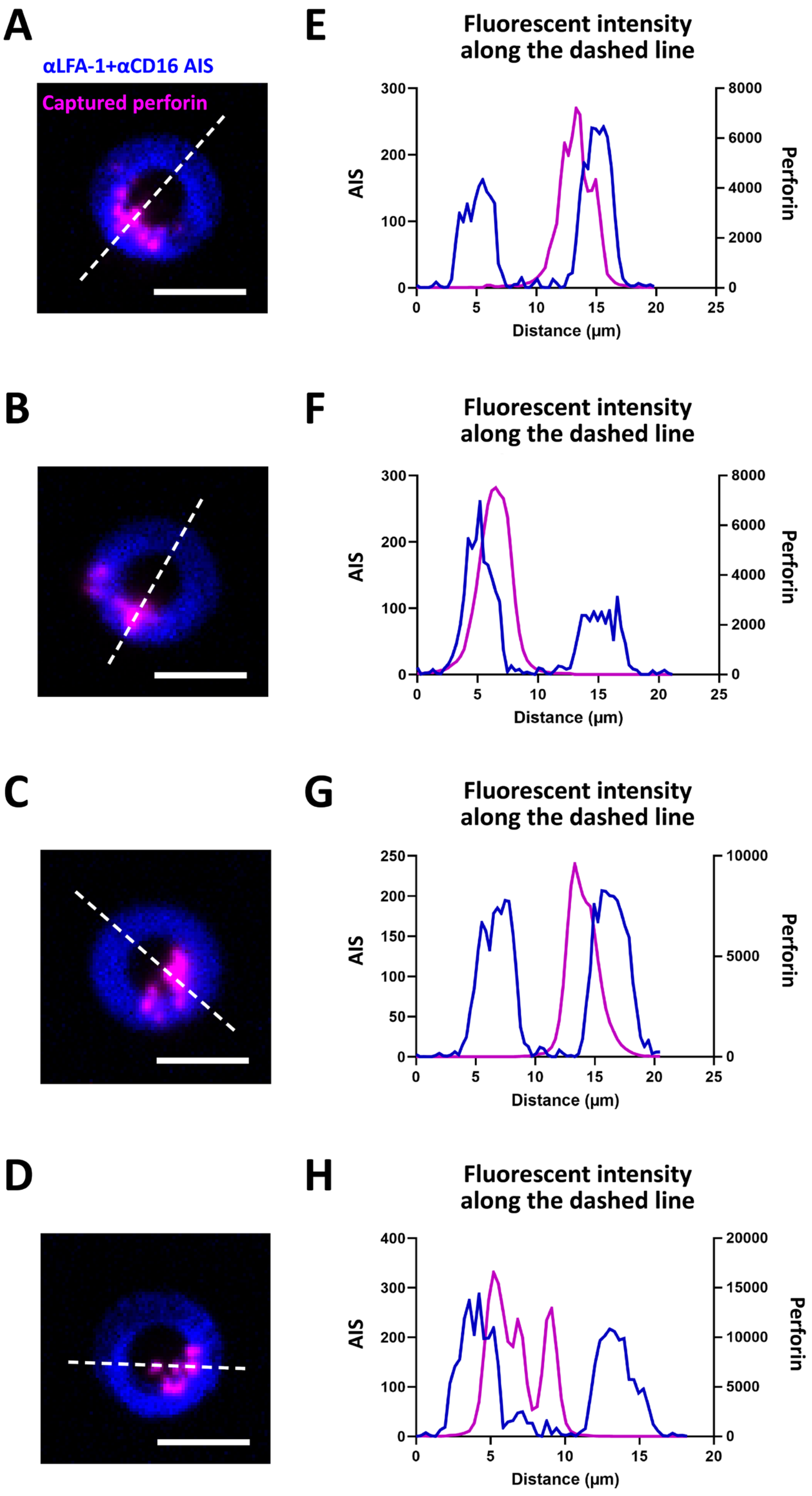


**Supporting Figure S5. Perforin can be captured across all regions of donut-shaped AIS.** αLFA-1 + αCD16 AIS + capture αPrf were printed on capture αPrf-coated substrates, allowing to capture released perforin from NK cells interacting with the AIS. To confirm capture in all regions of the AIS, large perforin clusters (often resulting from NK cells dying during the assay and releasing their entire perforin content) were selected from all experiments and the perforin distribution on the AIS was measured. **(A-D)** Fluorescent images of large captured perforin clusters, spanning over several AIS regions. **(E-H)** Fluorescent intensity profiles along the dashed lines indicated in (A-D). Perforin capture can be measured in all regions of the AIS, and no border effect can be observed at the inner edge of the AIS. Scale bars: 10µm.

**Supporting Movie 1: Single NK cell engaged in a prolonged, oscillating interaction with two αLFA-1 AIS simultaneously**

**Supporting Movie 2: NK cell forming a complete contact on αLFA-1 + αCD16 donut-shaped AIS**

**Supporting Movie 3: NK cell forming a partial contact on αLFA-1 + αCD16 donut-shaped AIS**

**Supporting Movie 4: Calcium activation upon synapse formation with αLFA-1 + αCD16 dot-shaped AIS**

**Supporting Movie 5: Calcium activation upon synapse formation with αLFA-1 + αCD16 donut-shaped AIS**

**Supporting Movie 6: Combined time-lapse imaging and capture of degranulated perforin, for an NK cell having formed a complete contact on αLFA-1 + αCD16 dot-shaped AIS.**

**Supporting Movie 7: Combined time-lapse imaging and capture of degranulated perforin, for an NK cell having formed a complete contact on αLFA-1 + αCD16 donut-shaped AIS.**
